# Neurostimulation Artifact Removal for Implantable Sensors Improves Signal Clarity and Decoding of Motor Volition

**DOI:** 10.1101/2022.09.07.506955

**Authors:** Eric J. Earley, Anton Berneving, Jan Zbinden, Max Ortiz-Catalan

## Abstract

As the demand for prosthetic limbs with reliable and multi-functional control increases, recent advances in myoelectric pattern recognition and implanted sensors have proven considerably advantageous. Additionally, sensory feedback from the prosthesis can be achieved via stimulation of the residual nerves, enabling closed-loop control over the prosthesis. However, this stimulation can cause interfering artifacts in the electromyographic (EMG) signals which deteriorate the reliability and function of the prosthesis. Here, we implement two real-time stimulation artifact removal algorithms, Template Subtraction and ε-Normalized Least Mean Squares, and investigate their performance in offline and real-time myoelectric pattern recognition in two transhumeral amputees implanted with nerve cuff and EMG electrodes. We show that both algorithms are capable of significantly improving signal-to-noise ratio and offline pattern recognition accuracy of artifact-corrupted EMG signals. Furthermore, both algorithms improved real-time decoding of motor intention during active neurostimulation. Although these outcomes are dependent on the user-specific sensor locations and neurostimulation settings, they nonetheless represent progress towards bi-directional neuromusculoskeletal prostheses capable of multifunction control and simultaneous sensory feedback.

## 1 Introduction

In 2008, Ziegler-Graham *et al*. estimated that the number of people living with amputations in the US would more than double by the year 2050 (Ziegler-Graham *et al*., 2008). Furthermore, it is estimated that there are more than 1 million annual limb amputations globally (Advanced Aputee Solutions LLC, 2012). This poses a significant challenge as the demand for prosthetic devices with reliable and multi-functional control for intuitive use in daily life increases.

During the last decades, most of the development has lead in the direction of introducing powered prostheses where movement control is decoded from surface electromyogram (sEMG) using myoelectric pattern recognition (Hudgins, Parker and Scott, 1993; Englehart and Hudgins, 2003; Zheng, Crouch and Eggleston, 2021). Further work has also been performed to improve control resolution and reliability through Targeted Muscle Reinnervation (TMR), where nerves in the remaining limb are innervated into existing musculature to increase the number of electromyogram channels and improve prosthesis controllability (Kuiken *et al*., 2004).

However, using sEMG for prosthesis control comes with a multitude of problems as the signal quality is heavily dependent on environmental conditions and susceptible to motion artifacts and myoelectric crosstalk (Ortiz-Catalan, Håkansson and Brånemark, 2014). To remedy this, recent work extending the concept of bone-anchored (osseointegrated) prostheses to also include bi-directional electrical communication has allowed electrodes to be implanted and connected directly through the implant to the prosthesis. This has improved controllability and general prosthesis usability over classical myoelectric prostheses (Ortiz-Catalan, Håkansson and Brånemark, 2014; Mastinu *et al*., 2018; Ortiz-Catalan, Mastinu, Sassu, *et al*., 2020).

The bi-directional communication additionally allows neurostimulation to provide sensory feedback to the user (Ortiz-Catalan, Håkansson and Brånemark, 2014; Ortiz-Catalan, Mastinu, Greenspon, *et al*., 2020). By placing spiral cuff electrodes around nerves in the residual limb, somatosensory (touch) sensations can be elicited through neurostimulation. However, due to the nature of electrical signals, and the fact that the electrical stimulation pulses are often larger in amplitude compared to the underlying EMG signal, the stimulation pulses can also be picked up by the nearby EMG electrodes. This creates unwanted artifacts in the activation patterns used to detect the user’s intent and leads to reduced pattern recognition performance and robustness (Hartmann *et al*., 2015).

The problem of stimulation artifacts (SAs) is not only present in the area of prosthesis control, but also applies to any system involving closed-loop neuromodulation (Zhou, Johnson and Muller, 2018). Literature on the topic of general artifact removal exists, but research involving prosthesis control applications is lacking. Moreover, the existing literature mostly focuses on removing electrocardiogram (ECG) artifacts from the EMG signals (Marque *et al*., 2005; Zhou, Lock and Kuiken, 2007) and do not consider the artifacts caused by neurostimulation for providing tactile feedback. Therefore, there is a need to develop and test methods for real-time stimulation artifact removal (SAR) using implanted EMG electrodes.

In this paper, we implement two real-time stimulation artifact removal algorithms, Template Subtraction and ε-Normalized Least Mean Squares, and investigate their performance in offline and real-time myoelectric pattern recognition with two transhumeral amputees implanted with nerve cuff and EMG electrodes. Using offline analysis, we show that both algorithms are capable of significantly improving signal-to-noise ratio and pattern recognition accuracy of artifact-corrupted EMG signals. Furthermore, both algorithms improved real-time decoding during a Motion Test (Todd A. Kuiken *et al*., 2009; Ortiz-Catalan, Brånemark and Håkansson, 2013) performed during active neurostimulation.

## 2 Methods

This study was approved by the Swedish regional ethical committee in Gothenburg (Dnr: 769-12). All participants provided written informed consent prior to participation in the study.

### 2.1 Hardware

The embedded hardware was based on previous work (Mastinu *et al*., 2017). It comprises a TM4C123GH6PM 32-bit ARM Cortex-M4F main micro controller unit (MCU) with floating point unit clocked at 80 MHz (Texas Instruments, USA) and a secondary MSP430G2755 16-bit mixed signal MCU at 16 MHz (Texas Instruments, USA). The secondary MCU handles stimulation waveform generation and signal acquisition through a RHS2116 digital electrophysiology stimulator and amplifier chip (Intan Technologies, USA), capable of sampling from implanted electrodes and stimulating the extraneural spiral-cuff electrodes within the arm of a person with an e-OPRA implant system (Integrum, Sweden). During stimulation pulse generation, the secondary MCU ceases signal acquisition, effectively blanking the EMG signal to avoid saturating the EMG channels and quickly return the electrode potential to baseline after each stimulation pulse (Hartmann *et al*., 2015). This blanking removes most of the SA spike but leaves the longer lasting exponential tail still in the signal (O’Keeffe *et al*., 2001; Zhou, Johnson and Muller, 2018). However, by sampling the stimulation channel, a reference signal highly correlated with the stimulation artifact can be acquired for later use by SAR algorithms.

### 2.2 Algorithm Selection & Description

Two algorithms were selected with focus on ease of implementation, computational complexity, and ability to perform SAR in real-time during closed-loop control of a prosthetic hand.

#### 2.2.1 Template Subtraction

We implemented a template subtraction algorithm based on a first order infinite impulse response (IIR) filter, similar to the ones presented by Keller *et al*. (Keller and Popovic, 2001) and Azin *et al*. (Azin, Chiel and Mohseni, 2007). The algorithm was chosen due to its simplicity and recursive formulation, yielding a computationally efficient implementation for usage in real-time on an embedded device.

By averaging artifacts from previous stimulation pulses using multiple exponential filters with filter learning rate *α*, a representative template of length *N* is constructed which is subtracted from the original signal *x*(*t*) to yield an estimated artifact-free signal 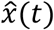 as illustrated in **Figure 1**.

**Figure 1.**
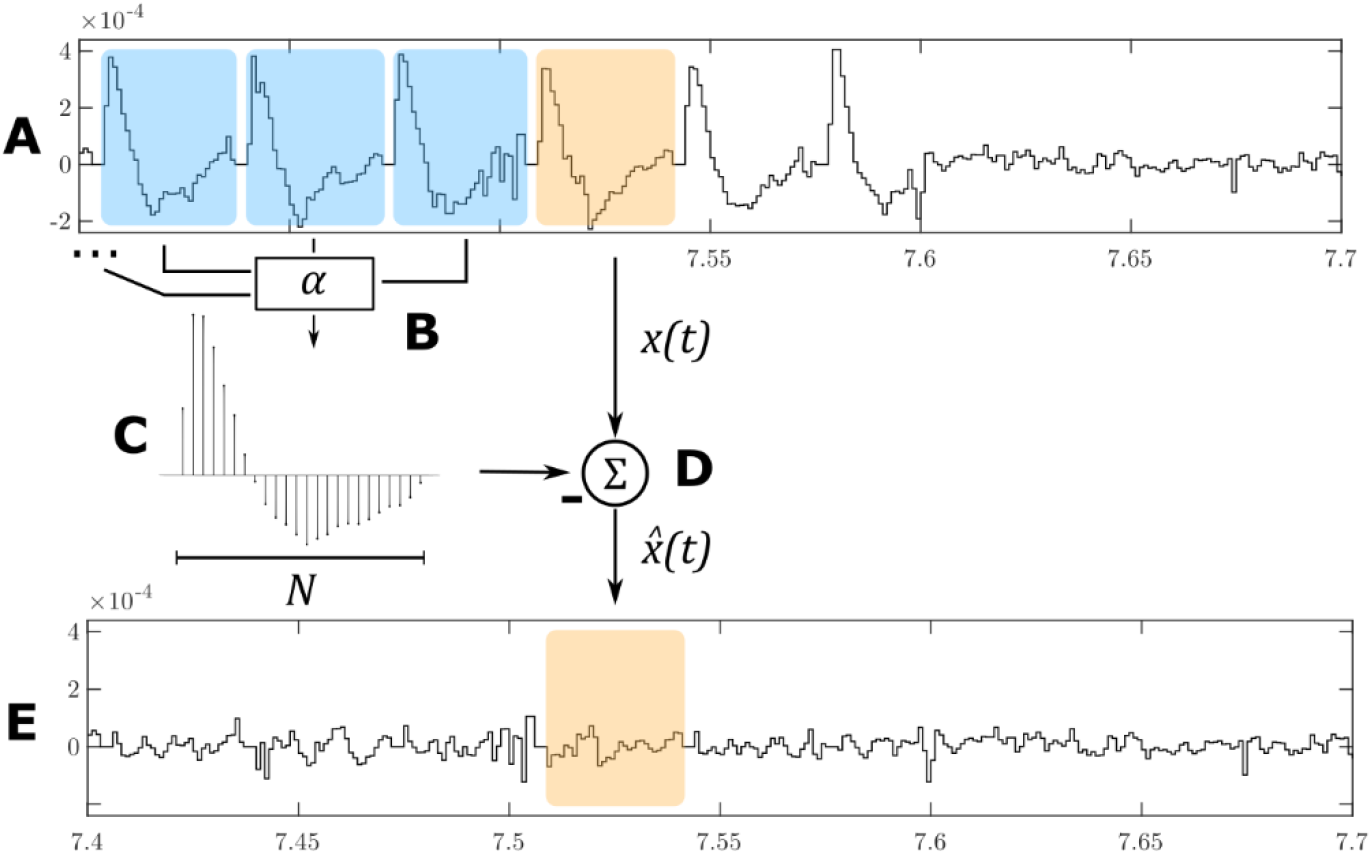
The Template Subtraction algorithm identifies stimulation artifacts from the original signal ***x***(***t***) (A, in blue) and averages them using exponential filters (B) to form an artifact template (C). This template is subtracted from a newly-identified artifact (D, in orange) to yield an estimated artifact-free signal 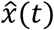 (E).

Let W_k_(*i*) represent the learned artifact at sample *i* = 1, …, *N* after the end of stimulation pulse *k*. Assuming that t_k_ is the sample index at which stimulation pulse *k* ends, i.e., when the stimulation artifact is present in the EMG signal, the recursive update is defined as:

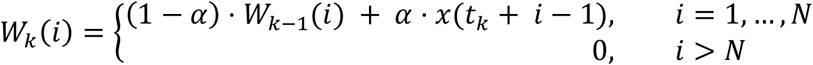

and the estimated artifact-free signal is then given by subtracting the template:

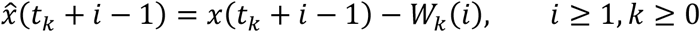

#### 2.2.2 ε-NLMS

We also implemented a variant of the common Least Mean Squares (LMS) adaptive filter, namely the ε-Normalized Least Mean Squares (ε-NLMS) filter. The ε-NLMS algorithm is an improved version of the standard LMS algorithm with a variable step size, yielding better performance for signals with intervals of larger and lower signal energy such as speech signals and overall faster convergence (Sayed, 2008). Due to the variable step size, the ε-NLMS algorithm is slightly more computationally expensive than the standard LMS algorithm, but has still been successfully used to remove neurostimulation artifacts in real-time (Basir-Kazeruni *et al*., 2017).

As the adaptive filter relies on a reference signal highly correlated with the SA (in our case, samples from the stimulation channel itself), the algorithm can adapt to varying artifact waveforms without requiring completely relearning of the weights. **Figure 2** illustrates the working principle of the adaptive filter.

**Figure 2.**
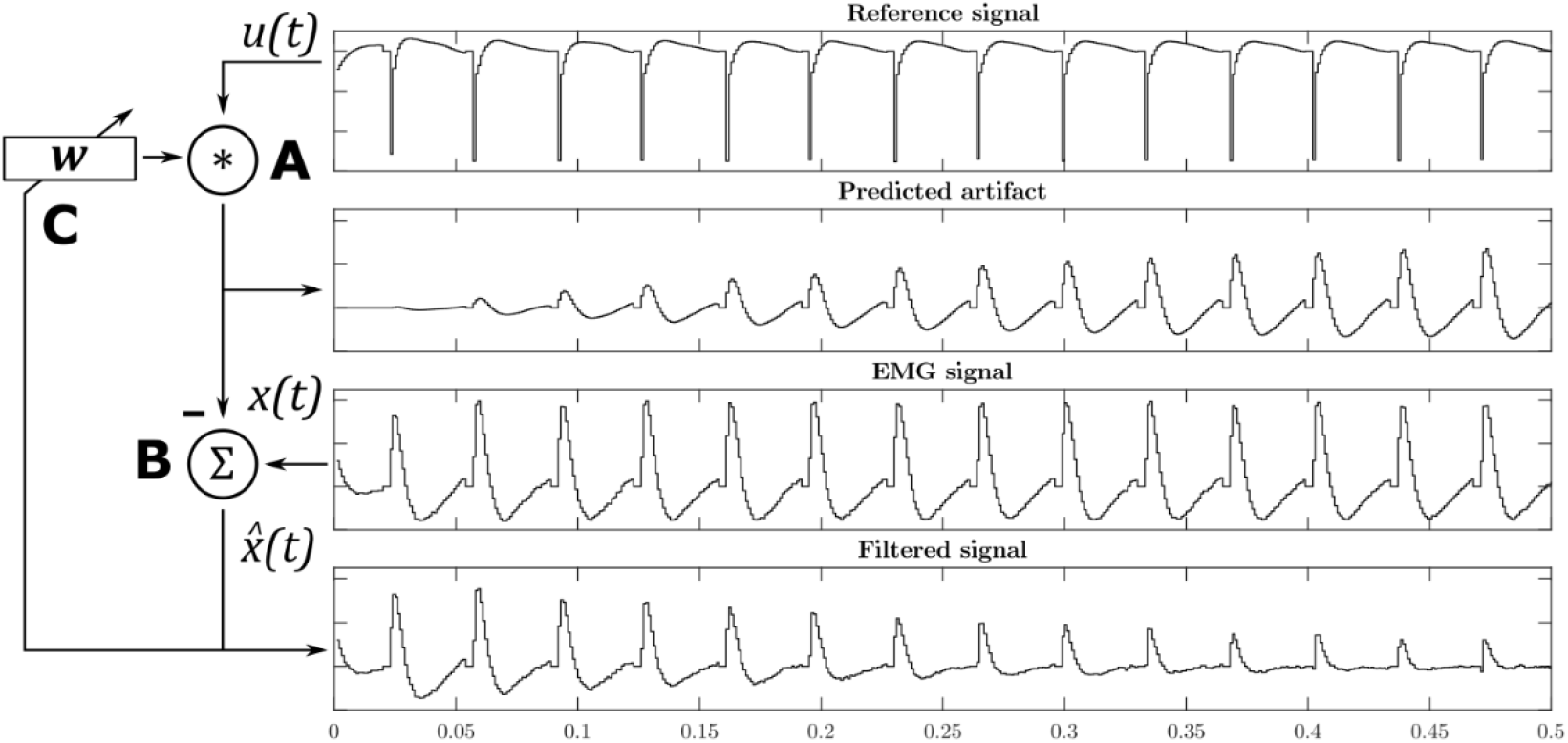
The ε-NLMS adaptive filter algorithm convolutes the reference signal ***u***(***t***) with filter weights ***w*** to predict the artifact waveform (A). The predicted artifact is subtracted from the original signal ***x***(***t***) to yield an estimated artifact-free signal 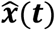 (B), which is then also used to update the filter weights after each sample (C).

Let *x*(*t*) be the original signal, ***u***(*t*) = [*u*(*t*) *u*(*t* − 1) … *u*(*t* − *N* − 1)] a vector containing the *N* previous samples from the reference *u*(*t*), and ***w***_*t*_ a weight vector of length *N* at sample *t*. The estimated artifact-free signal 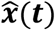 is then given as:

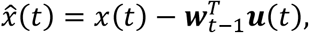

and the weights are updated according to the ε-NLMS update rule:

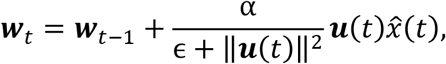

where *∈* > 0 is a small constant for avoiding division by zero, and *α* is a positive learning rate parameter.

### 2.3 Offline evaluation

An offline evaluation to identify optimal algorithm parameters and investigate the sensitivity to parameter variation was conducted with one participant with a transhumeral amputation and an osseointegrated e-OPRA implant system (Integrum AB) connected to a prosthetic arm as previously reported (Ortiz-Catalan, Mastinu, Sassu, *et al*., 2020). Signals were sampled at 1000 Hz from the four channels normally used to control *hand open, hand close, pronate*, and *supinate* functions using direct control. In addition, the stimulation channel was sampled as a reference for the ε-NLMS algorithm. Signals were processed with a 50 Hz notch filer to reduce electrical interference and passed through a second order digital high-pass filter at 20 Hz to remove signal bias. Square biphasic, asymmetric, stimulation pulses (Günter, Delbeke and Ortiz-Catalan, 2019) were applied at the stimulation frequency through the cuff electrode around the median nerve.

#### 2.3.1 Initial data collection

We performed an initial data collection to obtain EMG signals both with and without SAs for offline evaluation. During the experiment, the participant was asked to perform three tasks: (i) no movement, (ii) repeated *Hand Close* and *Hand Open*, or (iii) repeated *Pronation* and *Supination*. Signals in scenario (i) were recorded for 10 seconds, while scenarios (ii) and (iii) were recorded for 15 seconds each. During the first half of each recording, stimulation was applied in the form of multiple successive pulses, where four sets of suitable stimulation parameters were chosen based on the participant’s detection threshold – pulse amplitudes of 300 μA and 450 μA, pulse frequencies of 30 Hz and 50 Hz, and a pulse width of 150 μs. In total, 12 recordings were obtained containing both artifact-free and artifact-contaminated EMG signals.

#### 2.3.2 Algorithm parameter selection

To select the values of hyperparameters *N* and *α* for each algorithm, we devised an optimization scheme based on semi-synthetic signals. By using the data collected during scenario (i) and superimposing pure stimulation artifacts *a*(*t*) from scenario (ii) and (iii) onto the artifact-free EMG signals *x*(*t*), the Root Mean Square Error (RMSE) between the true artifact-free signal *x*(*t*) and the estimated artifact-free signal 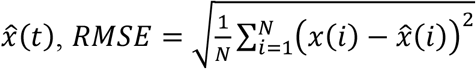, can be used as the optimization objective (Liang and Lin, 2002; Li, Chen and Yang, 2019). This approach has been used in previous work for evaluating SAR algorithm performance (Liang and Lin, 2002; De Clercq *et al*., 2006; Waddell *et al*., 2009). Due to the large variance in signal amplitude, the parameters were optimized for each channel individually and selected based on the median optimal RMSE of each of the 32 semi-synthetic signal combinations.

#### 2.3.3 Sensitivity analysis

Since selecting the algorithm hyperparameters is a complicated task when no existing signals are available for use in the above-described optimization scheme, we investigated the change in algorithm performance with respect to small deviations from the optimal parameters for *N* and *α*.

Using the same set of semi-synthetic signals as the optimization procedure, the algorithms’ hyperparameters were varied before applying them to the signals. However, using the RMSE between the true EMG signal *x*(*t*) and the estimated artifact-free signal 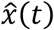 as a performance metric can be problematic for comparing performance between channels due to the large variation in signal amplitude. The RMSE is completely usable when optimizing algorithm performance, but as soon as comparing signals of different fundamental amplitudes, another metric is required. Additionally, the RMSE is difficult to interpret and relate in terms of absolute performance.

Therefore, a more general metric based on Signal-to-Noise Ratio (SNR), as employed in (Basir-Kazeruni *et al*., 2017), was used for evaluating the algorithms’ performance related to changes in the hyperparameters. The metric, hereby denoted as Δ*SNR* or SNR improvement, measures the change in SNR in decibels when applying the algorithm. It is not dependent on the absolute amplitude of the signals, but rather on the relative energy content of the true signal *x*(*t*), artifact *a*(*t*), and estimated artifact-free signal 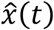, as

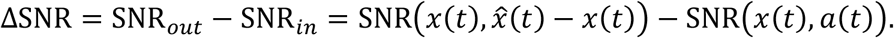

#### 2.3.4 Pattern recognition performance

To evaluate the impact of the algorithms on the pattern recognition performance, signals were manually labeled with both the intended movement and the presence or absence of simultaneous stimulation (**Figure 3**). Both SAR algorithms were applied to pre-recorded signals before separating the samples into time windows of 200 samples with 150 samples of overlap. In accordance with the findings by Hartmann et al. (Hartmann *et al*., 2015), blanked samples were removed from signals before extracting Mean Absolute Value (MAV), Zero Crossings (ZC), Slope Sign Changes (SSC), and Waveform Length (WL) from each window (Hudgins, Parker and Scott, 1993).

**Figure 3.**
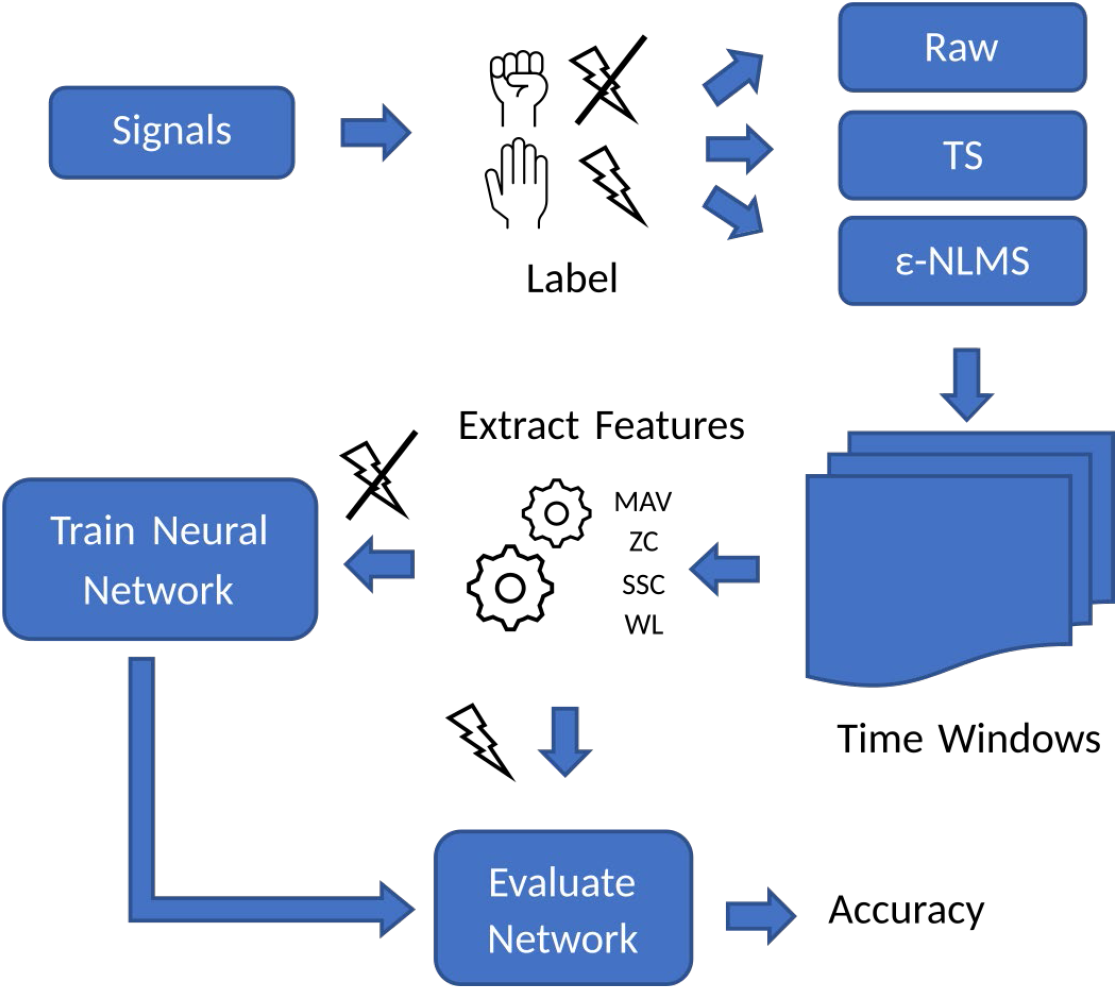
Overview of the process used for evaluating offline MPR performance of the two algorithms on signals containing stimulation artifacts.

The feature vectors from time windows without SAs were used to train a one-layer neural network using Rectified Linear Unit (ReLU) and softmax activation functions for predicting the intended prosthesis movement. The feature vectors used for training were randomly split into a 60% training set, 20% validation set and a 20% testing set, and the training set was augmented by adding additional 10db SNR noise to the feature vectors after being normalized. The network was trained and evaluated in MATLAB (MathWorks Inc., USA).

The remaining feature vectors, calculated on time windows containing SAs, both raw and processed by each algorithm, were fed to the trained network to evaluate the network and algorithm performance. The training and evaluation procedure was performed 50 times to account for the inherent randomness during data splitting and training.

### 2.4 Real-Time Evaluations

To evaluate the real-time implications for prosthesis controllability, *motion tests* were performed with two participants with trans-humeral amputation (T A Kuiken *et al*., 2009) as implemented in BioPatRec(Ortiz-Catalan, Brånemark and Håkansson, 2013). Both participants were home users of the osseointegrated e-OPRA implant.

First, a one layer neural network was trained on the standard features MAV, ZS, SSC, and WL (Hudgins, Parker and Scott, 1993) calculated from time windows at 500 Hz of length 200 ms, with 150 ms overlap on eight EMG channels to predict the prosthesis movements *Hand Open, Hand Close, Pronate, Supinate*, and *Rest*. No stimulation was active during the training and thus no SAs were included in the training data.

A series of Motion Tests were performed, where the participant was instructed to perform each movement multiple times in randomized order and hold the correct movement for at least 1 second within a time limit of 5 seconds while measuring the time to completion. First, a test without stimulation was performed to assess the baseline controllability of the prosthesis. Then, stimulation as a train of successive stimulation pulses at 20 Hz was enabled during the test; stimulation amplitude and pulse width were selected such that control of the prosthesis was impaired, but limited control of the prosthesis was still possible. Finally, the two algorithms were enabled on all signal channels and two additional motion tests (one for each algorithm) were performed. The TS algorithm was implemented with a learning rate *α* = 0.06 and a template length *N* = 25, and the ε-NLMS was implemented with a learning rate *α* = 0.035 and a filter length *N* = 25. The order of SAR algorithms was randomized.

### 2.5 Statistical analysis

For the offline evaluation, t-tests were performed to assess if there was a significant difference in classification accuracy for signals containing SAs and after applying the TS and ε-NLMS algorithms. Corrections for multiple comparisons were made using Holm-Bonferroni corrections.

For the Motion Test, the Wilcoxon rank sum test was performed to assess if there was a significant difference in completion rate, completion time, classification accuracy, and selection time between movements with and without SAs, and between the artifact-corrupted trials and the trials with SAR enabled. Corrections for multiple comparisons were made using Holm-Bonferroni corrections.

## 3 Results

### 3.1 Offline evaluation

Using the initial data collected, we performed a series of offline evaluations of the two algorithms to investigate their sensitivity to tuning of hyperparameters and effect on MPR classification accuracy.

#### 3.1.1 Sensitivity analysis

##### 3.1.1.1 Learning rate

As can be seen in **Figure 4**, larger learning rates *α* generally led to more variability in the *ΔSNR* for both algorithms, indicating that a larger learning rate improves SNR more for some semi-synthetic signals while causing a reduced improvement for others. For some parameter values, the *ΔSNR* also yielded negative changes, meaning that the algorithms induced more artifact noise into the signal than what was effectively removed. This was especially prominent for ε-NLMS, which displayed an overall larger performance variation.

**Figure 4.**
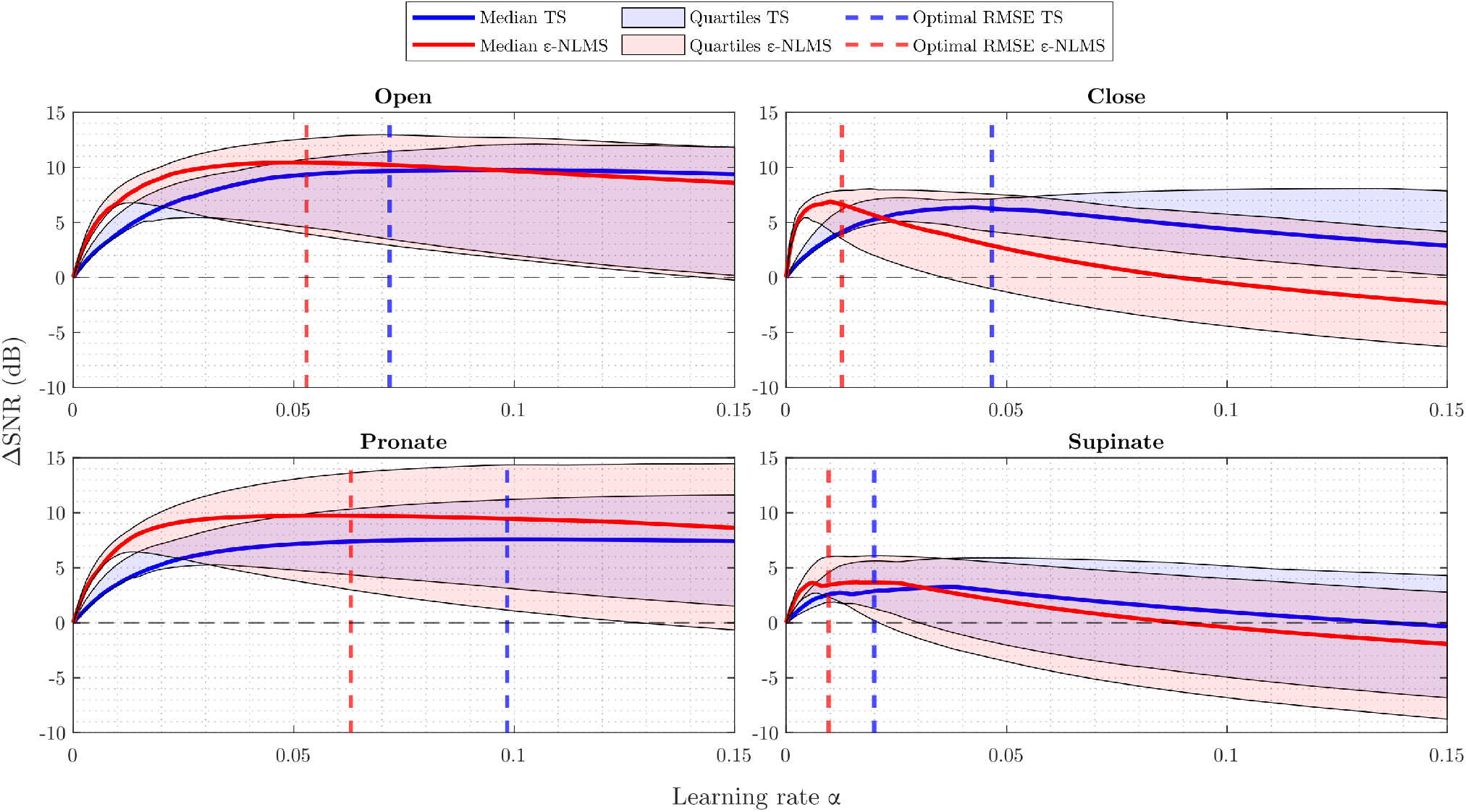
**Δ*SNR*** increased with an increasing learning rate ***α*** to a point, after which further increases in the learning rate worsened the change. The sensitivity of both TS (blue) and ε-NLMS (red) showed similar trends. Higher learning rates ***α*** led to increased variability (shaded regions), but the ***α*** at the optimal median RMSE between true and estimated signals (dashed lines) provided a good balance between median performance and variability.

By comparing the *ΔSNR* curves with the dashed lines representing the optimal parameter value for minimizing the median RMSE, it is evident that both performance metrics (RMSE and *ΔSNR*) were seemingly well correlated. The optimal parameters (dashed lines) generally provided a sound trade-off between the median and lower quartile SNR improvement. For example, consider the performance of the TS algorithm (blue) for the supinate channel (lower right) in Figure 4 – by purely optimizing median *ΔSNR*, the optimal parameter would be *α* = 0.04 which would yield a larger *ΔSNR* variability and even cause the lower quartile to lie below zero. Instead, the RMSE optima provided similar median and upper quartile *ΔSNR* performance, but with a considerably higher lower quartile performance.

In summary, an increased learning rate *α* led to increased performance variability and optimizing for RMSE provided a good balance between median performance and variability. Furthermore, an excessive learning rate caused negative *ΔSNR* improvement, indicating that *α* needs to be tuned sufficiently low for correct functionality, especially for ε-NLMS.

##### 3.1.1.2 Template/filter length

Considering the sensitivity when the template/filter length *N* was varied in **Figure 5**, the choice of *N* does not seem to be as crucial for determining the algorithms’ performance as the learning rate *α*.

**Figure 5.**
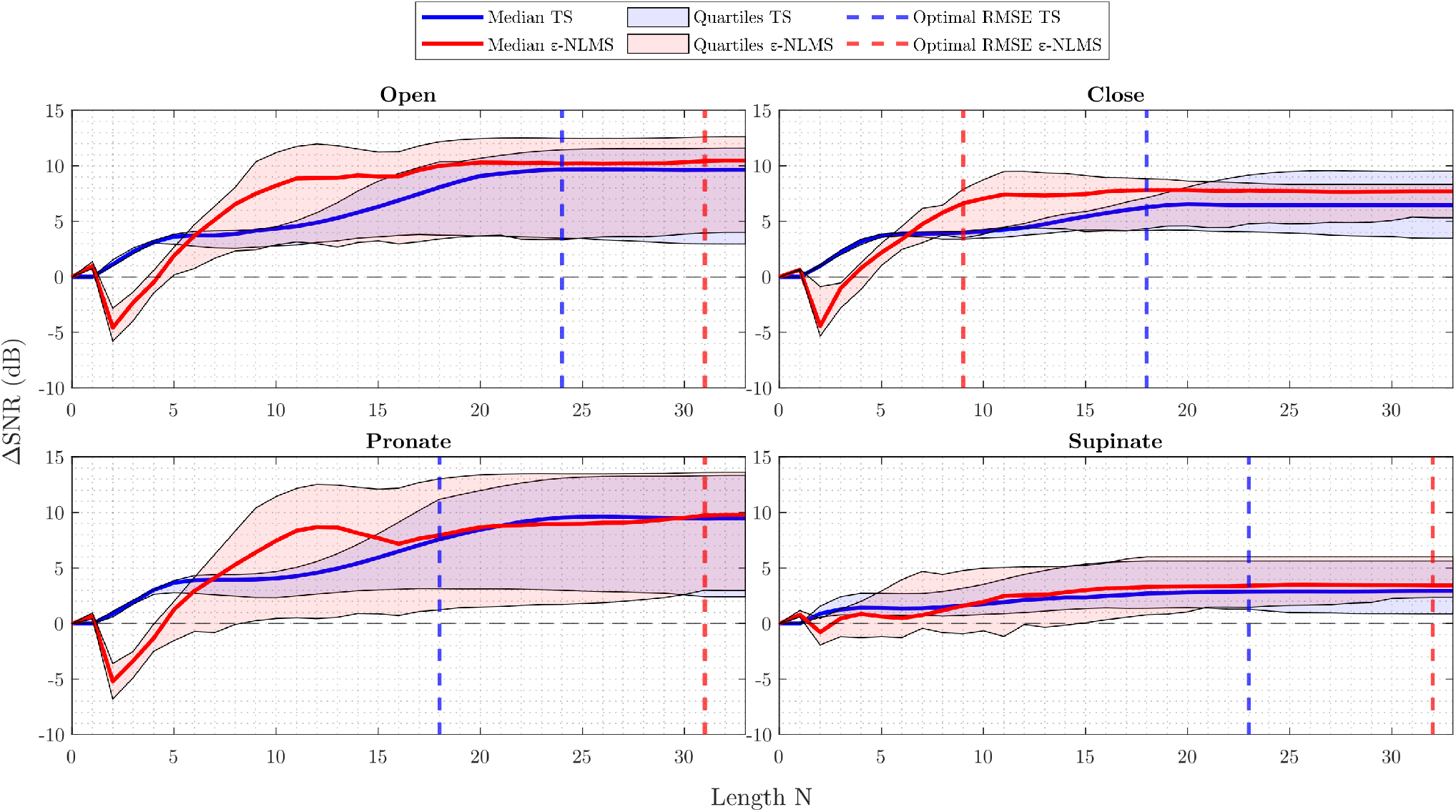
**Δ*SNR*** generally improved with an increasing length ***N*** for TS (blue), however ε-NLMS (red) required a minimum length be met before improvement was seen. Variability of the outcome (shaded region) did not appear to relate to the length ***N***, nor did the opimal median RMSE between true and estimated signals (dashed lines) consistently align with the maximum **Δ*SNR***.

One may observe that after a certain length the performance stayed constant. This is a logical behavior, considering that the number of samples between each stimulation pulse is dependent on the sampling and stimulation frequency. In the offline evaluation, the semi-synthetic signals were sampled at 1000 Hz and contained stimulation pulses generated at 30 and 50 Hz, leading to a maximum of roughly 33 samples between each pulse. It is therefore understandable that the algorithm performance stayed constant once *N* ≥ 30 as both algorithms are reset to start over again once a new pulse is detected.

Another interesting observation is that TS behaved similarly across all channels. When the template length was increased from zero, the SNR improvement was immediate and continued to increase when the template was extended. Considering ε-NLMS, however, a too short filter (*N* ≤ 7 in this case) decreased the SNR between the residual artifact and the true EMG signal when applying the algorithm, rather than removing the stimulation artifact. It thus seems important that the filter length is chosen long enough to correctly reduce the artifact signal power.

In contrast to the learning rate, both algorithms’ performance consistently improved as the template/filter length *N* increased. However, ε-NLMS required a sufficient length before any SNR improvement was noticed while TS improved the SNR immediately when increasing the length from zero. This indicates that ε-NLMS may require more care when manually tuning the filter length. Optimizing for RMSE seemed to provide a reasonable trade-off between performance and variability, although the observation was not as evident as for the learning rate.

#### 3.1.2 Offline Pattern Recognition

Using the constructed semi-synthetic signals containing artifacts, we evaluated the effect of stimulation on the MPR performance. In total, *n* = 100 training and evaluation iterations were performed to account for the inherent randomness of the training data split and network initialization. The total accuracy, defined as the portion of the predictions that were correct, decreased significantly on signals containing raw SAs (*p* < 0.001) and was significantly improved after applying either of TS or ε-NLMS (*p* < 0.001), see **Figure 6**. The accuracy did not reach the same levels as on the artifact-free test signals (*p* < 0.001) although TS performed better than ε-NLMS (*p* < 0.001).

**Figure 6.**
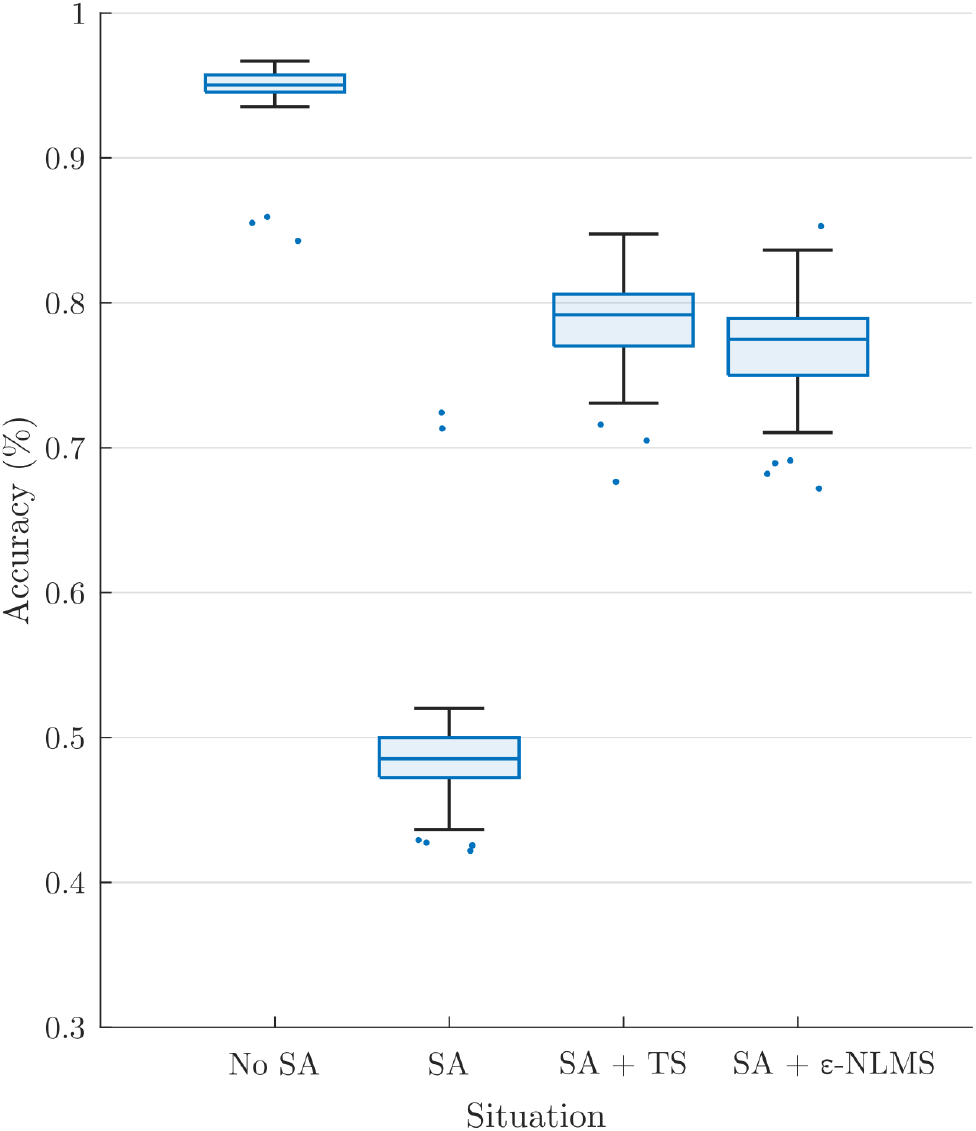
Offline pattern recognition accuracy was significantly reduced for signals containing SAs but both TS and ε-NLMS significantly restored the PR accuracy, although without reaching the same level as for the test set of artifact-free signals.

### 3.2 Motion Tests

The Motion Test was used to determine the effect of the two artifact removal algorithms on real-time myoelectric pattern recognition. Participants were generally able to complete movements when no stimulation was provided (96% completion rate), movements were significantly affected during stimulation (20% completion rate, *p* < 0.001, **Figure 7a**). This completion rate was significantly improved by using either TS (56%, *p* = 0.029) or ε-NLMS (52%, *p* = 0.041).

**Figure 7.**
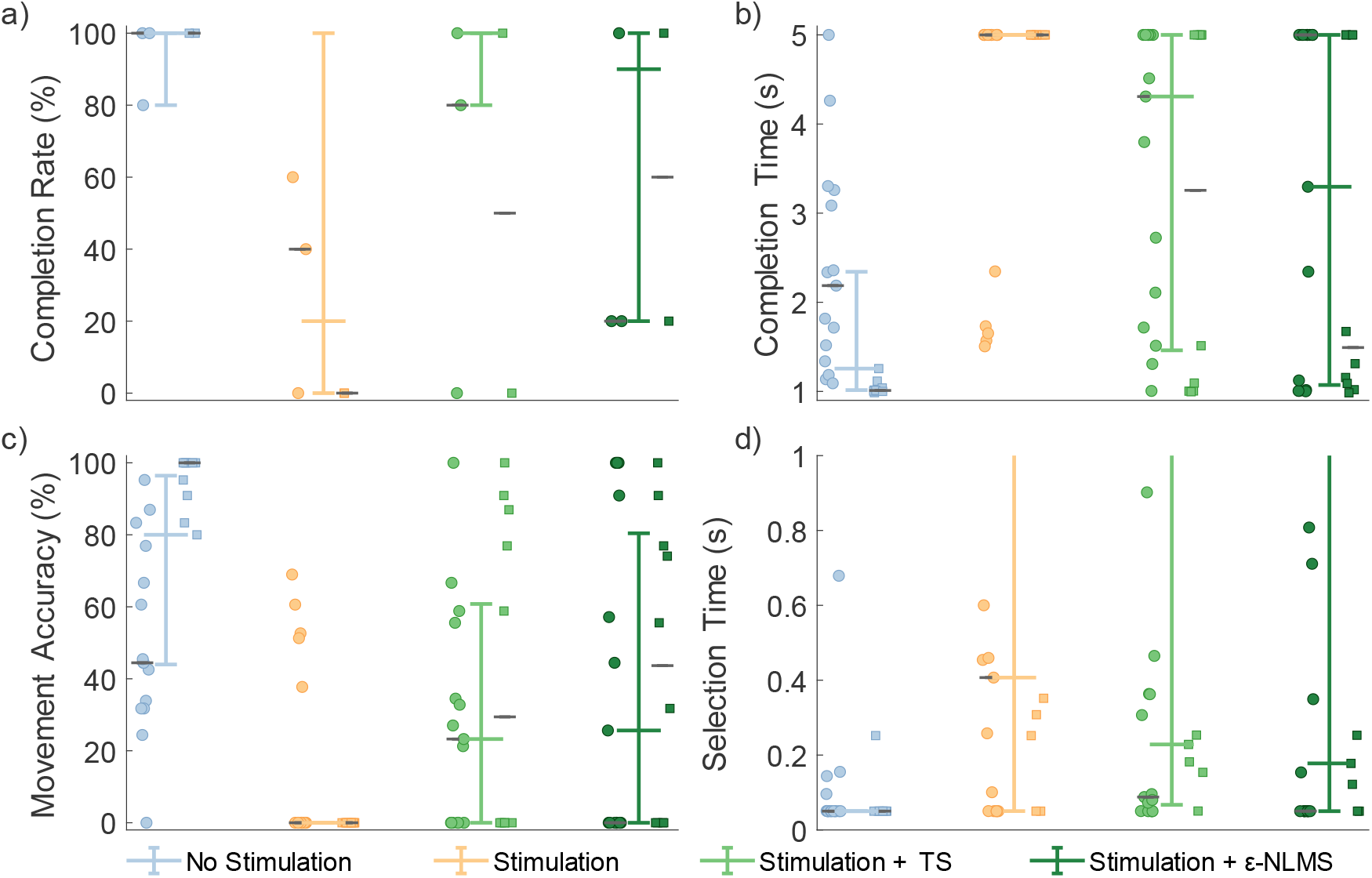
Real-time Motion Test outcomes were deteriorated by neurostimulation, but performance was recovered when using artifact removal algorithms. Template Subtraction resulted in a significantly improved completion rate (a), and ε-NLMS significantly reduced the movement completion time (b) and increased the movement accuracy (c). Neither algorithm had a significant effect on selection time (d). Colored whisker plots represent pooled median and interquartile range. Circle and square points represent data from the two participants, and the gray horizontal lines represent their individual medians.

Additional outcome measures from the Motion Test also provide insight into how the artifact removal algorithms affected myoelectric decoding. The completion time describes the amount of time taken to perform the defined movement for 1 second, within a 5 second trial. Completion time showed generally good performance without stimulation (median [IQR]: 1.26s [1.01, 2.34]), which significantly increased during neurostimulation (5s [5,5], *p* < 0.001, **Figure 7b**). Completion time was significantly recovered when using TS (4.31s [1.46, 5], *p* = 0.017) and ε-NLMS (3.30s [1.07, 5], *p* = 0.020). The movement accuracy indicated that the high rate of correct predictions without feedback (80.0% [44.0%, 96.4%]) similarly diminished with active sensory feedback (0% [0,0], *p* < 0.001, **Figure 7c**). Again similarly, this performance was significantly restored with TS (23.3% [0, 60.8], *p* = 0.021) and ε-NLMS (25.6% [0, 80.4], *p* = 0.030). Selection time did not show the same trend; while uncorrupted selection time (0.05s [0.05,0.05]) was still significantly hindered by the application of neurostimulation (0.41s [0.05, 5], *p* < 0.001), performance was not significantly different when using TS (0.23s [0.07, 1.93], *p* = 0.890) or ε-NLMS (0.18s [0.05, 2.09], *p* = 0.907, **Figure 7d**).

Taken together, these results suggest that, for movements that are significantly impacted by neurostimulation artifacts, the use of Template Subtraction and ε-NLMS artifact removal algorithms can help to restore some of the lost performance by making affected movements easier and quicker to achieve.

## 4 Discussion/Conclusion

In this paper, we implemented and evaluated two algorithms, Template Subtraction and ε-NLMS, to remove artifacts caused by neurostimulation used for somatosensory feedback. With the aim to restore control to myoelectric prosthesis users while neurostimulation for sensory feedback is provided, we evaluated the algorithms’ sensitivity to hyperparameters and their effect on offline and online pattern recognition performance.

Considering the algorithms’ sensitivity to variations in the two hyperparameters *α* and *N*, the results suggest that an increase in the learning rate *α* generally leads to higher variability with increased performance for some semi-synthetic signals and worsened performance for others. The increased variability additionally led to a negative *ΔSNR* caused by more noise being introduced in the signal than what was removed in terms of artifacts. On the other hand, optimizing the RMSE seemed to provide a good balance between median performance and performance variability, suggesting that the RMSE is a useful optimization metric for automatic selection of hyperparameters.

Neither algorithm was negatively affected by an increase in the length parameter *N*, but the improvement was constant once the inter-pulse period was reached. Additionally, ε-NLMS required a sufficient filter length (*N* ⪆ 7) to yield improvements in SNR, while TS provided a consistent improvement starting from *N* = 1. This indicates that the ε-NLMS adaptive filter is dependent on enough previous samples on the reference channel to be able to predict the artifact waveform correctly. Furthermore, TS seems to require a larger *N* to reach the same performance as ε-NLMS. Given the two algorithms’ different approaches to predicting the artifact, this behavior is expected. The TS approach, of constructing a template for each of the *N* samples following the stimulation pulse, naturally requires a large enough *N* depending on the length of the artifact in the time domain for enough effect. On the other hand, when basing the prediction on the *N* last samples, as in ε-NLMS, a lower but large enough *N* is sufficient to adequately predict the artifact.

When comparing the two algorithms in offline analyses, there is no clear evidence supporting that one is significantly better than the other. In the evaluation of the algorithms’ sensitivity to variations in the hyperparameters, both algorithms performed similarly in terms of achieved SNR improvement. ε-NLMS, however, seemed to require more care when selecting parameters since a too high learning rate *α* quickly degraded the lower quartile performance and a too low filter length *N* induced more noise in the signal, ultimately causing a negative SNR improvement. In this respect, TS, which had a larger range of values of *α* that provided sufficient performance and showed consistent performance increase for all values of *N*, gives the impression of being easier to tune manually. TS is additionally more computationally efficient than ε-NLMS due to its simpler recursive update formulation.

Regarding the real-time Motion Test outcomes, both TS and ε-NLMS showed to the movement completion rate, completion times, and accuracy, suggesting that either algorithm can be used to recover raw signals corrupted by SAs. However, it should also be noted that these outcomes are dependent on numerous other factors. Our offline investigations suggest that hyperparameter selection can greatly impact the performance of the SAR algorithms, and more so that the optimal hyperparameter values may differ substantially between channels. In the real-time tests, we opted to use the same hyperparameters for all channels, demonstrating a general-use scenario, however fine tuning and selection of hyperparameters may lead to further improved SAR effectiveness.

As is often the case with prosthetics research, user variability will also drastically impact the need for and benefit of using SAR algorithms. In this study, two participants with transhumeral neuromusculoskeletal prostheses tested our algorithms; despite the fact that the amputation level and technology was similar between our participants, differences in surgical reconstruction, electrode and nerve cuff placement, and prosthesis use meant that different muscles were used to control the same prosthesis movements, and depending on the proximity of the nerve cuffs to each electrode the relative strength of SAs could differ. Thus, stimulation parameters had been selected for each participant to achieve a balance of performance; if SAs were too small, control of the prosthesis would not be affected and SAR algorithms would demonstrate no benefit, and if SAs were too large, the raw signal would be unrecoverable with SAR and the control would remain compromised. The stimulation parameters selected for this study managed this balance, but as a result the outcomes of this study must be interpreted within this basis – outside of this balanced stimulation window, the benefit of these SAR algorithms is reduced.

The TS and ε-NLMS algorithms are both well suited to learn and counteract static artifacts, which makes it particularly suited for stimulation paradigms such as discrete event-driven sensory feedback (Cipriani *et al*., 2014), which have consistent and repeatable stimulation patterns. However, most sensory feedback research sets the stimulation intensity proportional to the feedback measurement (typically grip force). Both TS and ε-NLMS can adapt to changing stimulation artifacts, however they will always lag behind a proportional feedback scheme. While these algorithms may still reduce the SA, they may under- or overcorrect and continue to leave residual artifacts in the signals. SAR methods which take the current stimulation parameters into account may be able to circumvent this issue, which is an area we plan to investigate in a future study.

Overall, when possible, it is best for implanted electrodes to be configured in such a way as to minimize the potential for SAs. Evaluation methods such as cross-channel impedance measurement may be used to identify configurations with a higher likelihood for SAs (Earley, Mastinu and Ortiz-Catalan, 2022), and the use of biopolar electrodes may help to reduce the likelihood of cross-talk. However, monopolar configurations can allow for a greater number of unique muscle sources for a given number of electrodes, which is of particular importance for implanted electrodes. In these cases, SAR algorithms may be able to make up the difference and improve the control of prosthetic limbs while simultaneously permitting sensory feedback, allowing for bidirectional and closed-loop control of prostheses which serve to improve the independence and quality of life of people with amputations.

## 5 Conflict of Interest

The authors declare that the research was conducted in the absence of any commercial or financial relationships that could be construed as a potential conflict of interest. MOC has been consultant for Integrum AB, and the rest of the authors declare no conflict of interest.

## 6 Author Contributions

AB, EJE, and JZ developed and implemented the algorithms. EJE and AB developed the experimental protocols. AB, EJE, and JZ ran the experiments. EJE and AB analyzed the data. AB and EJE prepared the figures. EJE and AB drafted the manuscript. MO-C envisioned the study and obtained funding for this project. All authors read and approved the final manuscript.

## 7 Funding

This work was supported by the Promobilia Foundation, the IngaBritt and Arne Lundbergs Foundation, and the Swedish Research Council (Vetenskapsrådet).

## Acknowledgments

We would like to acknowledge and thank our participants for their contributions and feedback to this study.

## 10 Data Availability Statement

The datasets generated for this study can be found at the Open Science Framework (Earley, Berneving and Ortiz-Catalan, 2022).

## Notes

https://osf.io/y473e/

